# Acute Milk-Protein Intake Enhances Pupil-Linked Executive Function and Esports Performance During Prolonged Play

**DOI:** 10.64898/2026.03.23.713804

**Authors:** Takashi Matsui, Shion Takahashi, Daisuke Funabashi, Chisato Ohba, Kentaro Nakamura

**Affiliations:** Institute of Health and Sport Sciences, University of Tsukuba, 1-1-1 Tennoudai, Tsukuba, Ibaraki 305-8574, Japan; R&D Center for Sport Innovation, University of Tsukuba, 1-1-1 Tennoudai, Tsukuba, Ibaraki 305-8574, Japan; R&D Division, Meiji Co., Ltd., Tokyo 192-0919, Japan

**Keywords:** esports, cognitive fatigue, milk protein, executive function, pupillometry, physiological markers, nutrition

## Abstract

Prolonged esports play induces cognitive fatigue that is not fully captured by subjective awareness, motivating practical, non-stimulant nutritional strategies supported by objective physiological markers. We here tested whether acute milk protein intake attenuates fatigue-related physiological responses during prolonged esports play and supports subjective state, executive control, and in-game performance. In a randomized, single-blind (assessor-blind), energy-matched controlled crossover study, 15 healthy young adults with esports experience completed two sessions in which they consumed either a milk protein drink or an energy-matched apple juice control before a 3-h virtual soccer task. Physiological measures included pupillometry during gameplay, salivary cortisol, continuous interstitial glucose monitoring, and heart rate. Subjective ratings (VAS) and executive function (flanker task) were assessed across post-ingestion time points, and in-game performance metrics were aggregated within hourly gameplay blocks. Milk protein intake was associated with a coherent pattern of physiological advantages, including larger pupil diameter during gameplay, smoother interstitial glucose dynamics, and lower salivary cortisol, while heart rate showed time-dependent changes without a clear condition effect. These physiological changes co-occurred with higher enjoyment and lower hunger, improved flanker performance, and condition-dependent improvements in in-game performance, most notably higher shot success rate. Additionally, pupil diameter during gameplay was associated with inhibitory-control efficiency on the flanker task. These findings suggest that acute milk protein intake may serve as a practical, non-stimulant nutritional strategy to sustain physiological state and cognitive–behavioral performance during prolonged esports (virtual soccer) play.

**Highlights:**

– Prolonged esports play models modern digital cognitive activity and cognitive fatigue.
– Acute milk protein intake increases pupil diameter during prolonged esports play.
– Interstitial glucose dynamics are smoother and salivary cortisol is lower with milk protein.
– Enjoyment increases and hunger decreases during 3 h of virtual soccer play.
– Executive function and in-game performance improve, most notably shot success rate.

## 1. Introduction

Esports has rapidly evolved into a major global industry, supported by professional leagues, sponsorship, streaming platforms, and a growing ecosystem of training and performance-support services (Białecki et al., 2024; Hong, 2023). Alongside elite competition, a much larger population of recreational players now spends extended periods in cognitively demanding game environments, making esports a relevant setting for both high-performance optimization and broader public well-being (Hong, 2023).

Prolonged esports play can induce cognitive fatigue that may emerge before players fully recognize subjective tiredness, potentially compromising executive control and performance during critical moments (DiFrancisco-Donoghue et al., 2021). This dissociation motivates the use of objective physiological markers to track fatigue accumulation and evaluate countermeasures. Recent evidence using pupillometry indicates that extended esports play is accompanied by progressive pupil constriction that covaries with declines in cognitive performance, suggesting that pupillary responses can sensitively index cognitive fatigue during prolonged play (Matsui et al., 2024).

Physiological state changes during prolonged play are also plausibly linked to subjective experience and cognitive performance. Pupil dynamics have been linked to neuromodulatory systems relevant to cognitive control, including locus coeruleus–noradrenergic activity, supporting pupillometry as a window into task engagement and cognitive effort (Joshi et al., 2016; van der Wel & van Steenbergen, 2018). In addition, prolonged cognitive effort, including esports play, is accompanied by stress- and autonomic-related responses that may be reflected in markers such as salivary cortisol and heart rate (Hellhammer et al., 2009; Hjortskov et al., 2004; Kraemer et al., 2022), while metabolic dynamics such as glycemic fluctuations may influence appetite-related sensations and perceived energy during extended sessions (Arumugam et al., 2008). In particular, postprandial glucose dynamics, including glycaemic “dips,” have been linked to subsequent hunger and energy intake in healthy individuals, supporting the relevance of glycaemic stability to appetite-related sensations during prolonged tasks (Wyatt et al., 2021).

As a result, multi-domain measurement spanning physiology, subjective state, executive function, and in-game behavior is needed to characterize fatigue trajectories and to identify practical interventions that can be implemented during esports play. In this sense, esports can serve as a tractable model of prolonged digital cognitive activity, sharing key fatigue demands with other modern contexts such as computer-based work, training, or learning. However, empirically supported, implementable countermeasures for multi-hour esports sessions remain limited, and many studies focus on discrete sessions rather than sustained play in which fatigue accumulates (DiFrancisco-Donoghue et al., 2019; Sousa et al., 2020).

In this context, we use the term “smart nutrition” to refer to targeted, short-term nutritional interventions implemented immediately before or during esports play that aim to sustain cognitive–behavioral performance while minimizing potential health trade-offs (e.g., sleep disruption or excessive sugar load). Nutritional strategies may influence cognitive control partly by modulating the availability of amino acid precursors for neurotransmitters and neuromodulators relevant to executive function (Lieberman, 2003). Milk protein–based drinks have been reported to improve aspects of executive function in healthy young adults (Nouchi et al., 2023; Saito et al., 2018), suggesting potential utility for fatigue-prone esports contexts.

At the same time, players often rely on caffeine- and sugar-containing beverages to transiently restore alertness, yet repeated or excessive use raises concerns regarding sleep disruption and metabolic burden, motivating the evaluation of non-stimulant alternatives (Gardiner et al., 2023; Kennedy & Scholey, 2004; Malik et al., 2010; Nadeem et al., 2021). In addition, rapidly absorbed carbohydrates may induce pronounced glycaemic fluctuations, including postprandial “dips,” which have been linked to subsequent increases in hunger and variability in perceived energy, potentially undermining sustained cognitive performance during prolonged tasks (Wyatt et al., 2021). Emerging evidence, including a randomized crossover study reporting that a caffeine- and sugar-free beverage (sparkling water) attenuated fatigue-related pupil constriction during prolonged esports play, further supports the feasibility of in-game, non-stimulant strategies targeting cognitive fatigue (Takahashi et al., 2026). However, whether acute milk protein intake can sustain executive function and in-game performance during prolonged esports play while also attenuating fatigue-related physiological signatures such as pupil constriction has not been directly tested in crossover settings.

Accordingly, we hypothesized that acute milk protein intake attenuates cognitive fatigue-related pupil constriction and improves subjective state, executive control, and in-game performance. To test this hypothesis, we conducted a randomized, single-blind, energy-matched controlled crossover study in healthy young adults with esports experience, assessing physiological measures (pupillometry, salivary cortisol, CGM-derived interstitial glucose, and heart rate), as well as VAS ratings, flanker performance, and in-game metrics.

## 2. Materials and Methods

**2.1 Study design**

We conducted a randomized, single-blind, energy-matched controlled crossover study to examine the acute effects of a milk protein–based drink on cognitive fatigue and performance during prolonged esports play. Each participant completed two experimental sessions under two beverage conditions (milk protein drink vs. control beverage) in a counterbalanced order. In each session, participants consumed the assigned beverage and then performed an extended virtual soccer (esports) task, during which physiological, subjective, cognitive, and behavioral outcomes were repeatedly assessed. Pupillometry was used as an objective physiological marker of fatigue-related responses during gameplay, alongside other physiological measures (salivary cortisol, interstitial glucose, and heart rate), subjective ratings, executive function performance, and in-game performance indices derived from match statistics.

### 2.2 Participants

Fifteen healthy young adults with esports experience participated in this study. Participant characteristics are summarized in Table 1. The sample had a mean (± SD) age of 24.2 ± 4.9 years and included 15 men and 0 woman. Mean body weight was 63.4 ± 11.1 kg, height was 1.7 ± 0.1 m, and body mass index (BMI) was 22.2 ± 3.1 kg/m². Participants reported an average playing career in their proficient title of 10.5 ± 10.3 months and an in-game rank corresponding to the top 66.8 ± 41.2 percentile (0–100 scale, from highest to lowest rank). Self-reported esports playing time was 78.0 ± 71.0 min/day on weekdays and 120.0 ± 102.0 min/day on weekends.

**Table 1.** Participant characteristics of playing lifestyles.

|  |  | n = 15 |
| --- | --- | --- |
| Age | Years | 24.2 ± 4.9 |
| Sex | Male/female | 15 / 0 |
| Weight | kg | 63.4 ± 11.1 |
| Height | m | 1.7 ± 0.1 |
| BMI | kg/m<sup>2</sup> | 22.2 ± 3.1 |
| Playing career of proficient title | months | 10.5 ± 10.3 |
| In-game rank | top percentile | 66.8 ± 41.2 |
| Weekday esports playing time | min/day | 78.0 ± 71.0 |
| Weekend esports playing time | min/day | 120.0 ± 102.0 |
| Sleeping time | min/day | 420.0 ± 42.4 |
| Vigorous-intensity activity time | min/day | 64.4 ± 87.6 |
| Moderate-intensity activity time | min/day | 100.0 ± 119.1 |
| Walking time | min/day | 31.7 ± 23.1 |
| Exercise time | min/day | 40.0 ± 76.5 |
| Sitting time | min/day | 400.0 ± 256.3 |

Data are shown as mean ± SD. Sex data are presented as the number of male and female participants.
The in-game rank data show percentages from the highest to the lowest rank, ranging from 0 to 100.

We conducted an a priori power analysis using G*Power 3.1.9.7, assuming an effect size of Cohen’s f = 0.27 based on a previous study that demonstrated cognitive fatigue in casual esports players (Matsui et al., 2024). The analysis indicated that a total of 20 participants would be sufficient to detect a significant within–between interaction in a repeated-measures ANOVA at an alpha level of .05 and a statistical power (1 – β) of .80, with two conditions, four repeated measurements, a correlation among repeated measures of .50, and a non-sphericity correction ε of 1.

Eligibility criteria included age between 18 and 35 years, capacity to understand and sign informed consent, normal or corrected-to-normal vision sufficient for video game play, normal color vision, and availability to attend the experiment in person. Exclusion criteria were a history of neurological disease, metabolic disease, or sleep disorders, current use of medication, and smoking.

Participants were recruited via a laboratory website and social media announcements. All participants provided written informed consent prior to participation. The study protocol was approved by the Research Ethics Committee of the Faculty of Health and Sport Sciences, University of Tsukuba (approval No. 021–112), and conducted in accordance with the Declaration of Helsinki. A washout period of at least 3 days separated the two experimental visits in the randomized crossover design. Participant flow, including enrollment, randomization, and crossover allocation, is summarized in Supplementary figure 1.

To standardize pre-visit conditions, participants were instructed to abstain from caffeine on the test day and from alcohol intake and strenuous exercise on the day before each visit. Participants were asked to maintain their usual sleep habits. They were also instructed to finish their last meal by 21:00 on the evening before each visit and to arrive at the laboratory on the test day without consuming any energy-containing foods or beverages.

### 2.3 Test beverages and blinding

Two energy-matched beverages were used: a milk protein drink and an energy-matched control beverage. The milk protein drink (SAVAS MILK PROTEIN Fat 0, Milk Flavor (Meiji Co., Ltd., Tokyo, Japan) provided 102 kcal per serving (200 mL) and contained 15.0 g protein, 10.4 g carbohydrate, 0 g fat, and 95.4 mg sodium. In addition, the control beverage (apple juice) provided 102 kcal per serving (237.2 mL) and contained 25.3 g carbohydrate, 0 g protein, 0 g fat, and 11.9 mg sodium. Nutrient composition and serving volumes are summarized in Table 2.

**Table 2.** Nutrient composition of the test beverages (per serving).

| **Nutrient** | **Control beverage**<br>**(Apple juice)** | **Milk protein**<br>**beverage** |
| --- | --- | --- |
| Serving volume (mL) | 237.2 | 200.0 |
| Energy (kcal) | 102.0 | 102.0 |
| Protein (g) | 0.0 | 15.0 |
| Fat (g) | 0.0 | 0.0 |
| Carbohydrate (g) | 25.3 | 10.4 |
| Sodium (mg) | 11.9 | 95.4 |

The study adopted a single-blind procedure. Beverage conditions were coded by a third party not involved in data collection or analysis, and investigators responsible for outcome assessment were blinded to condition assignment until completion of the primary analyses. To minimize sensory cues and maintain blinding for investigators, beverages were served in identical opaque containers, and participants and outcome assessors were blinded to condition assignment. Each participant consumed one full serving of the assigned beverage at the beginning of each experimental session..

### 2.4 Experimental procedure

The overall experimental timeline and assessment schedule are illustrated in Figure 1. Each participant completed two laboratory visits under the two beverage conditions in a randomized crossover design, separated by a washout period of at least 3 days. All visits started between 08:30 and 09:00. Indoor environmental conditions were standardized: illuminance was maintained at 250–300 lux (both at the center of the room and at the participant’s eye level), and room temperature was kept at 22 ± 1.0°C.

**Figure 1.**
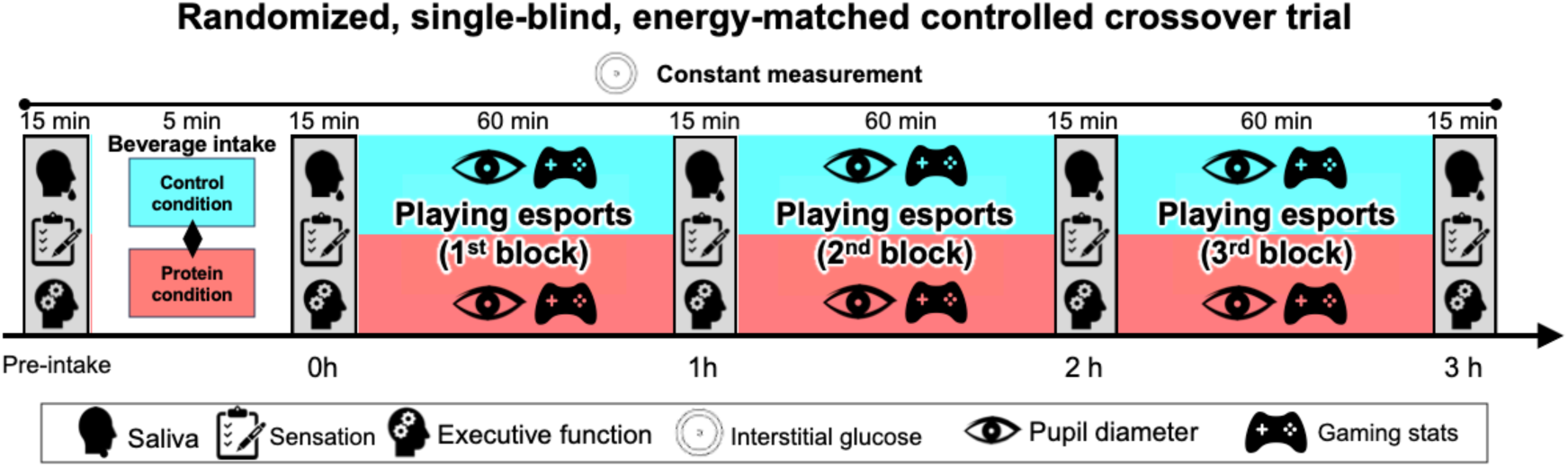
Experimental design and assessment timeline. Schematic overview of the randomized, single-blind (assessor-blind), energy-matched controlled crossover trial. After arrival, participants rested quietly in a seated position for 10 min before baseline assessments (Pre-intake). Participants then consumed the assigned beverage (milk protein drink or energy-matched apple juice control), and post-ingestion assessments were conducted at 0 h prior to initiating gameplay. Participants subsequently performed a continuous 3-h virtual soccer task (eFootball), divided into three 1-h gameplay blocks. At 1 h, 2 h, and 3 h after ingestion, gameplay was paused for a standardized 15-min break to complete assessments. Measures included salivary cortisol, subjective ratings (VAS), executive function (flanker task), interstitial glucose (CGM), heart rate, pupillometry (during gameplay), and in-game performance metrics.

Upon arrival, compliance with pre-visit restrictions was confirmed. Participants then rested quietly in a seated position for 10 min without conversation or smartphone use, after which baseline assessments were obtained prior to beverage ingestion (Pre-intake). Participants then consumed one full serving of the assigned beverage (milk protein drink or apple juice control) from an identical opaque container. Immediately after ingestion (0 h), post-ingestion assessments were completed before initiating the esports task.

#### Esports task

The esports task consisted of playing a virtual football game from the *eFootball* series (Konami Digital Entertainment, Tokyo, Japan) continuously for 3 h. Participants played offline matches against a computer-controlled opponent throughout each session. Game settings were standardized and held constant across conditions and visits, including match mode, opponent difficulty, and camera view. Match duration was set to allow multiple full matches to be completed within each 1-h block. Participants were seated at a fixed distance from the display and used a standard game controller. They were instructed to play as competitively and consistently as possible throughout the session, maintaining their usual tactics and formations and avoiding deliberate changes in play style between visits. Verbal encouragement was minimized to reduce external influences on motivation.

#### Timing of assessments

Physiological measures (including salivary cortisol, CGM-derived interstitial glucose, and heart rate) and pupillometry were collected according to the schedules described below; subjective ratings (VAS) and executive-function testing were administered at Pre-intake and 0 h prior to gameplay. During the 3-h esports task, assessments were performed at approximately 1 h, 2 h, and 3 h after ingestion (+1 h, +2 h, +3 h). At each time point, gameplay was paused and assessments were conducted during a standardized 15-min break. In-game performance metrics were recorded from the post-match summary screen after each match and aggregated within each 1-h gameplay block (1–3 h), as described in Section 2.5.

### 2.5 Outcome measures

#### 2.5.1 Physiological measures

##### Pupillometry

Pupil diameter was recorded as an objective index of central arousal and cognitive load during esports play, following the protocol described by Matsui et al. (2024). Binocular pupil size was measured using an infrared eye tracker (Tobii Pro Nano; Tobii Technology, Sweden) continuously throughout the 3-h *eFootball* task. Ambient lighting was standardized to 250–300 lux at eye level, and monitor brightness settings were held constant across visits. Under these conditions, visual stimuli were expected to be comparable across participants, beverage conditions, and time points.

The eye tracker was mounted below the display and connected to a dedicated laptop positioned next to a 23.8-inch monitor (60 Hz, 1920 × 1080 pixels). Participants were seated approximately 60 cm from the screen. Before the first experimental session, a four-point calibration was performed using fixation targets presented at the four corners of the game screen. Calibration accuracy was verified in Tobii Pro Lab (Tobii Technology, Sweden) by visually confirming that the recorded gaze position matched the instructed fixation targets.

Binocular pupil diameter data were recorded at 60 Hz during gameplay. Pupil data were preprocessed in Tobii Pro Lab. Due to device limitations, blinks could not be explicitly detected. Short data losses of up to 80 ms were interpolated using the software’s built-in gap fill-in function. Noise reduction was performed using the built-in moving median filter (window size: 5 samples). The final pupil diameter time series was computed as the average of the left and right eyes. To focus on longer-term trends rather than second-to-second fluctuations, data were exported as 1-min averages, and hourly averages were subsequently calculated for each of the three 1-h gameplay blocks (1 h, 2 h, 3 h). Because pupil diameter could only be recorded during gameplay, baseline measurements and normalization relative to pre-task levels were not performed.

##### Salivary cortisol

Salivary cortisol was measured as a physiological marker of stress associated with prolonged esports play. Saliva samples were collected at Pre-intake and 0 h prior to gameplay, and at approximately +1 h to +3 h during the standardized assessment breaks between gameplay blocks. For the present study, post-ingestion cortisol outcomes from 0 h to +3 h were considered the main time window, whereas Pre-intake values were treated as supplementary data.

At each time point, participants provided saliva by spitting directly into sterile collection tubes through a straw. Approximately 2 mL of saliva were collected at each time point and immediately stored at −80°C. To remove mucus, samples were centrifuged at 1500 × g for 20 min; the supernatant was aliquoted and stored at −80°C until assay. Cortisol concentrations were determined using a commercially available enzyme-linked immunosorbent assay kit (Salivary Cortisol ELISA kit; Salimetrics, LLC, State College, PA, USA) according to the manufacturer’s instructions. All samples were assayed in duplicate, and absorbance was read with a microplate reader (Varioskan LUX Multimode Microplate Reader; Thermo Fisher Scientific, Waltham, MA, USA). Cortisol concentrations were calculated from standard curves, and duplicate values were averaged for statistical analyses.

##### Interstitial glucose (continuous glucose monitoring)

Interstitial glucose concentration was monitored continuously using a professional continuous glucose monitoring system (FreeStyle Libre Pro; Abbott, Japan) attached to the upper arm. The sensor was inserted and initialized according to the manufacturer’s instructions during the familiarization visit (2–7 days before the first experimental session).

Glucose data were recorded continuously across each experimental visit, including the Pre-intake assessment period and the subsequent 3-h esports task. Glucose values were exported at 15-min intervals. For analyses aligned with the gameplay structure, 15-min glucose values were averaged within each 1-h block after ingestion (1 h, 2 h, 3 h) to yield participant-level hourly glucose measures. Pre-intake glucose was defined as the CGM value immediately preceding beverage ingestion (i.e., the most recent 15-min sample, which overlapped the seated rest period).

##### Heart rate

Heart rate was recorded continuously throughout each session using an optical heart rate sensor (Polar Verity Sense; Polar Electro, Finland). Data were sampled at 1 Hz and averaged into 1-min epochs. For analyses aligned with the gameplay structure, 1-min values were further averaged within each 1-h gameplay block (1 h, 2 h, 3 h) to yield participant-level hourly heart-rate measures. Heart rate during Pre-intake was calculated as the mean over the 10-min seated rest period prior to beverage ingestion, and heart rate at 0 h was calculated as the mean over the 15-min post-ingestion assessment period prior to initiating gameplay.

#### 2.5.2 Subjective ratings (VAS)

Subjective states were assessed using 100-mm visual analogue scales (VAS). Participants rated six items: enjoyment and fatigue (primary subjective outcomes), and sleepiness, hunger, craving for sweets, and thirst (secondary subjective outcomes). For each item, participants marked their current state on a horizontal 0–100 mm line, with anchors indicating “not at all” (0 mm) and “extremely” (100 mm).

VAS ratings were collected at Pre-intake and immediately post-ingestion (0 h) prior to gameplay, and at approximately 1 h, 2 h, and 3 h after ingestion (+1 h, +2 h, +3 h) during the standardized breaks within the 3-h esports task. For the present study, post-ingestion ratings from 0 h to +3 h were considered the main time window for VAS analyses, whereas Pre-intake ratings were treated as supplementary.

#### 2.5.3 Executive function task

Executive function was assessed using a computerized flanker task implemented in PsychoPy3 (v2021.1.2) (Peirce et al., 2019), with identical task parameters to those used in our previous study that successfully detected cognitive fatigue induced by prolonged virtual football play (Matsui et al., 2024). The task was executed on a desktop PC (Windows 10; Intel Core i7-11700 CPU; NVIDIA GeForce RTX 3070 Ti GPU; 32 GB RAM) and presented on a 23.8-inch LCD monitor (60 Hz refresh rate; 1920 × 1080 pixels).

On each trial, a central target arrow was flanked by two arrows on each side, pointing either in the same direction (congruent trials) or the opposite direction (incongruent trials). Participants were instructed to respond as quickly and accurately as possible to the direction of the central arrow by pressing the corresponding key on a response device. For each assessment, stimuli were presented in a fully randomized order with 32 congruent and 32 incongruent trials (64 trials total). Each trial consisted of a fixation cross (250 ms), stimulus presentation (up to 2000 ms), and a blank screen (1000 ms). The stimulus disappeared upon keypress and was followed by the blank screen until the next trial.

To reduce learning effects, participants completed three practice sets (192 trials total) across two occasions: during a preliminary training visit and immediately before the first baseline (Pre-intake) administration of the flanker task (Rappaport et al., 2025). Reaction time (RT) and accuracy were recorded for all trials. Trials with RTs <200 ms or >1500 ms were excluded as anticipations or lapses. Mean RT was calculated separately for congruent and incongruent trials. Flanker interference was quantified as the RT difference between incongruent and congruent trials (incongruent minus congruent). For accuracy, the primary metric was the percentage of correct responses in incongruent trials.

The flanker task was administered at Pre-intake, 0 h, and at approximately +1 h to +3 h during each visit. For the present study, post-ingestion outcomes from 0 h to +3 h were considered the main time window for executive-function analyses, whereas Pre-intake outcomes were treated as supplementary.

#### 2.5.4 In-game performance metrics

In-game performance was quantified using match statistics from the eFootball post-match summary screen. For each match within a session, the experimenter manually recorded the following six variables: goals scored, goals conceded, shots on target, shot success rate, total passes, and saves.

For analytical clarity, performance metrics were grouped a priori into three domains. Offensive performance included goals scored, shots on target, and shot success rate, reflecting the effectiveness of attacking actions. Defensive performance included goals conceded and saves, reflecting defensive control and error prevention. Total passes were treated as an intermediate metric reflecting overall play involvement rather than purely offensive or defensive performance.

For analysis, match-level values were averaged within each 1-h block of gameplay (1 h, 2 h, 3 h) to yield participant-level hourly performance measures (i.e., repeated measures across time). Because these statistics are only available during gameplay, in-game performance analyses were based on the 1–3 h gameplay blocks.

### 2.6 Statistical analysis

All statistical analyses were performed using GraphPad Prism 10 (GraphPad Software, San Diego, CA, USA). Statistical significance was set at *p* < 0.05 (two-tailed). Descriptive data are presented as mean ± SD unless otherwise stated.

For outcomes collected at multiple post-ingestion time points, analyses were performed using repeated-measures models with condition (milk protein vs. control), time, and their interaction (condition × time). The crossover sequence/order of conditions was randomized and recorded for each participant; period effects (visit 1 vs. visit 2) and sequence effects were evaluated where applicable for crossover analyses. The washout interval (≥3 days) was intended to minimize carryover.

To align with the hypothesized sequence from physiology to subjective and behavioral outcomes, primary analyses focused on post-ingestion measurements. For physiological, VAS, and executive-function outcomes collected at 0–3 h, condition × time models were applied to the 0–3 h time points, with Pre-intake values treated as supplementary and reported separately. For in-game performance and pupillometry, analyses were conducted on the 1–3 h gameplay blocks only because these measures are only defined during gameplay.

P-values were adjusted for multiple comparisons using the Holm–Bonferroni method when omnibus tests indicated significant main effects or interactions and post hoc pairwise comparisons were conducted as appropriate (e.g., between conditions at each time point and/or between time points within each condition). Where applicable, effect sizes and 95% confidence intervals were reported.

## 3. Results

### 3.1 Physiological measures

#### Pupillometry

Pupil diameter during gameplay (1-3h) decreased over time and differed between beverage conditions (Fig. 2A). A 2-way RM ANOVA revealed a significant main effect of time, *F*(2, 28) = 5.453, *p* = 0.0119, η²p = 0.331, and a significant main effect of condition, *F*(1, 14) = 18.81, *p* = 0.0012, η²p = 0.631, whereas the time × condition interaction was not significant, *F*(2, 28) = 0.3687, *p* = 0.698, η²p = 0.050. Across the three 1-h gameplay blocks, the protein condition showed a larger pupil diameter than the control condition (predicted mean difference = 0.263 mm, 95% CI [0.129, 0.396]).

**Figure 2.**
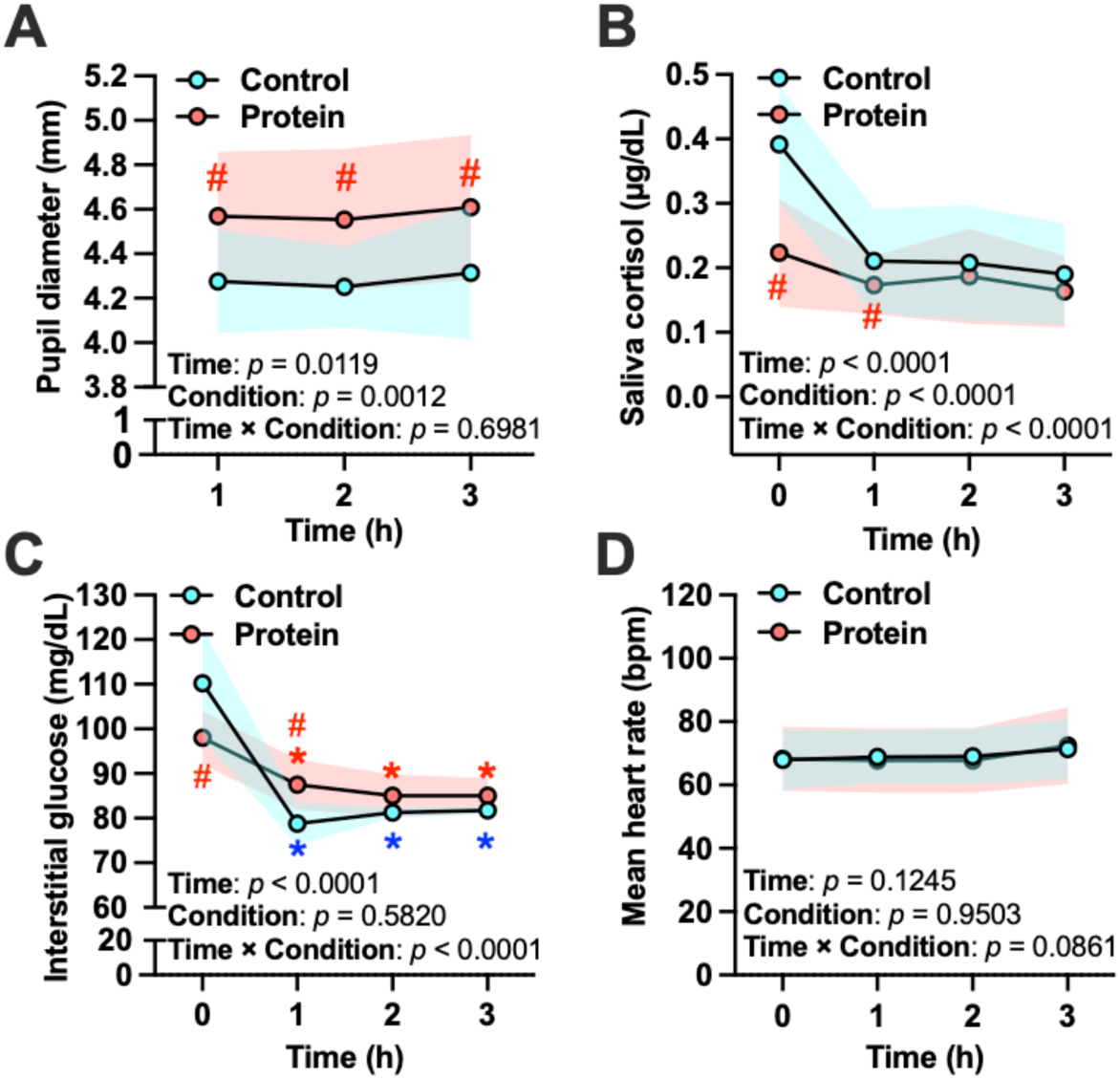
Physiological responses during prolonged esports play. Physiological measures under the milk protein and control conditions. (A) Pupil diameter during gameplay averaged within each 1-h block (1-3 h). (B) Salivary cortisol across post-ingestion time points (0–3 h). (C) Interstitial glucose measured by continuous glucose monitoring (CGM) across post-ingestion time points (0–3 h). (D) Heart rate across post-ingestion time points (0–3 h). Data are shown as mean ± SD. Results of the condition × time repeated-measures ANOVA are shown in the lower-left corner of each graph (Time, Condition, and Interaction). Where indicated, post-hoc comparisons were performed with Holm–Bonferroni correction. *: *p* < 0.05 vs. 0 h (a stable pre-ingestion reference time point) within the same condition. #: *p* < 0.05 vs. control condition at the indicated time point.

#### Salivary cortisol

A 2-way repeated-measures ANOVA revealed significant main effects of time (0-3h), *F*(3, 42) = 18.25, *p* < 0.0001, η²p = 0.566, and condition, *F*(1, 14) = 32.65, *p* < 0.0001, η²p = 0.700, as well as a significant time × condition interaction, *F*(3, 42) = 19.19, *p* < 0.0001, η²p = 0.578. Overall, cortisol concentrations were lower in the milk protein condition than in the control condition (difference between means = 0.063 µg/dL, 95% CI [0.039, 0.087]) (Fig. 2B).

#### Interstitial glucose (CGM)

A 2-way RM ANOVA revealed a significant main effect of time (0-3h), *F*(3, 42) = 73.05, *p* < 0.0001, η²p = 0.839, and a significant time × condition interaction, *F*(3, 42) = 19.01, *p* < 0.0001, η²p = 0.760, whereas the main effect of condition was not significant, *F*(1, 14) = 0.318, *p* = 0.582, η²p = 0.022. Mean glucose levels did not differ between conditions when averaged across time (difference between predicted means = −0.80 mg/dL, 95% CI [−3.83, 2.23]), but the temporal glucose profile differed markedly between conditions (Fig. 2C).

#### Heart rate

There was no significant main effect of time (0-3h), *F*(3, 42) = 4.180, *p* = 0.1245, η²p = 0.455. Neither the main effect of condition, *F*(1, 14) = 0.004, *p* = 0.950, η²p = 0.001, nor the time × condition interaction, *F*(3, 42) = 2.852, *p* = 0.086, η²p = 0.437, reached statistical significance (Fig. 2D).

#### Supplementary results

In supplementary analyses focusing on the baseline-to-immediate post-ingestion interval (Pre-intake to 0 h), milk protein intake was accompanied by lower salivary cortisol and a distinct interstitial glucose response, whereas mean heart rate did not differ between conditions (Supplementary Fig. 2). These findings indicate that milk protein intake acutely modulates stress- and glucose-related physiology without detectable changes in cardiac load prior to gameplay.

### 3.2 Subjective ratings (VAS)

Subjective ratings across post-ingestion time points (0–3 h) are shown in Figure 3. For enjoyment, there was a significant main effect of time, *F*(3, 42) = 12.95, *p* < 0.0001, η²p = 0.541, and a significant main effect of condition, *F*(1, 14) = 5.262, *p* = 0.0425, η²p = 0.324, whereas the time × condition interaction did not reach significance, *F*(3, 42) = 2.658, *p* = 0.0669, η²p = 0.216. Overall, enjoyment was higher in the milk protein condition than in the control condition (difference between predicted means = 6.92 mm, 95% CI [0.28, 13.56]) (Fig. 3A).

**Figure 3.**
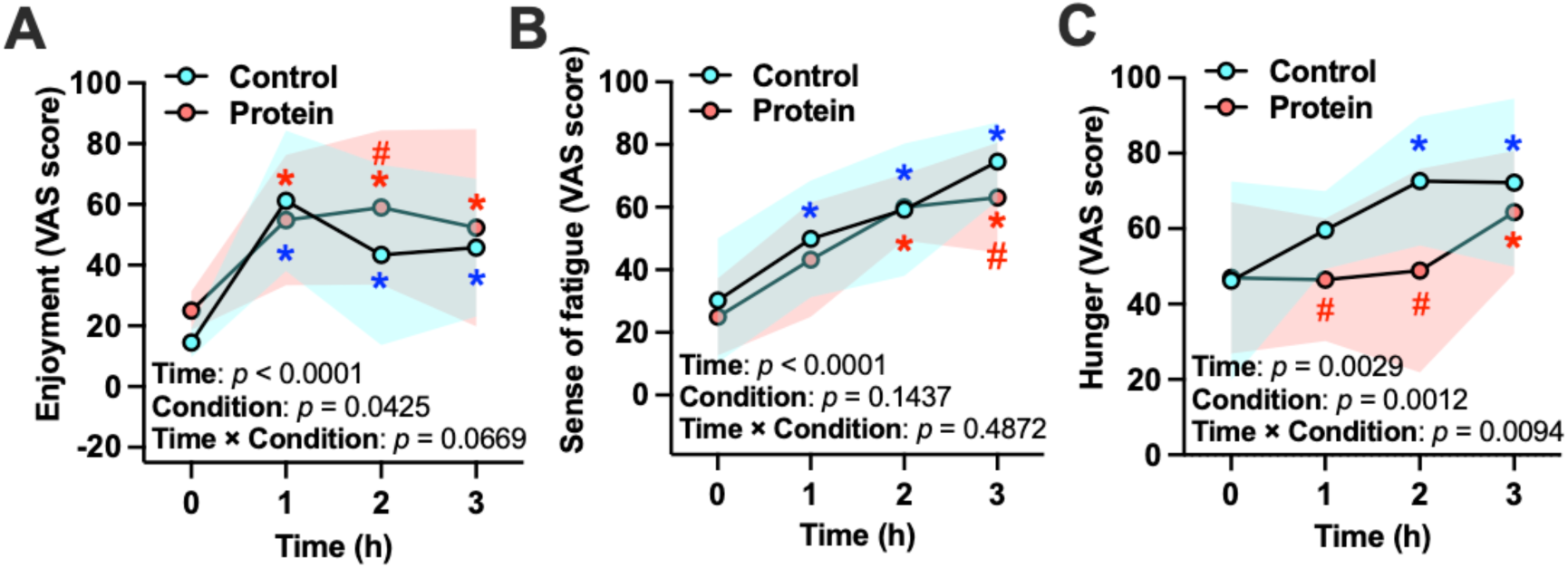
Subjective ratings during esports play. Visual analogue scale (VAS; 0–100 mm) ratings across post-ingestion time points (0–3 h) under the milk protein and control conditions. (A) Enjoyment. (B) Sense of fatigue. (C) Hunger. Values are shown as mean ± SD. The shaded bands represent the SD for the control and protein conditions. Results of the condition × time repeated-measures ANOVA are shown in the lower-left corner of each graph (Time, Condition, and Interaction). Where indicated, post-hoc comparisons were performed with Holm–Bonferroni correction. \**p* < 0.05 vs. 0 h within the same condition. #*p* < 0.05 vs. control condition at the indicated time point.

For fatigue, there was a significant main effect of time, *F*(3, 42) = 35.65, *p* < 0.0001, η²p = 0.764. The main effect of condition, *F*(1, 14) = 2.479, *p* = 0.1437, η²p = 0.184, and the time × condition interaction, *F*(3, 42) = 0.829, *p* = 0.4872, η²p = 0.070, were not significant (Fig. 3B).

For hunger, significant main effects of time, *F*(3, 42) = 5.966, *p* = 0.0029, η²p = 0.399, and condition, *F*(1, 14) = 21.90, *p* = 0.0012, η²p = 0.709, were observed, along with a significant time × condition interaction, *F*(3, 42) = 4.669, *p* = 0.0094, η²p = 0.342. Overall, hunger ratings were lower in the milk protein condition than in the control condition (difference between means = 11.01 mm, 95% CI [5.69, 16.33]) (Fig. 3C).

Analyses for craving for sweets, thirst, and sleepiness are reported as supplementary outcomes (Supplementary Fig. 3). Ratings for craving for sweets showed a significant main effect of time and interaction across 0–3 h (Time: F(3, 42) = 3.175, p = 0.0449, η²p = 0.388; Time × Condition: F(3, 42) = 2.555, p = 0.0443, η²p = 0.338; n = 6), whereas a main effect of condition, F(1, 14) = 0.656, p = 0.4548, η²p = 0.116, was not significant. For thirst, significant main effects of time and condition were observed (Time: F(3, 42) = 5.164, p = 0.0049, η²p = 0.319; Condition: F(1, 14) = 6.958, p = 0.0231, η²p = 0.387), whereas the time × condition interaction was not significant (F(3, 42) = 1.725, p = 0.1809, η²p = 0.136). For sleepiness, significant main effects of time and condition were observed (Time: F(3, 42) = 8.095, p = 0.0004, η²p = 0.424; Condition: F(1, 14) = 5.197, p = 0.0436, η²p = 0.321), whereas the time × condition interaction was not significant (F(3, 42) = 2.380, p = 0.0873, η²p = 0.178).

In supplementary analyses examining subjective responses from baseline to immediately after ingestion (Pre-intake to 0 h), hunger ratings decreased rapidly in both conditions, indicating a prompt reduction in perceived hunger irrespective of beverage type (Supplementary Fig. 4). In contrast, enjoyment, fatigue, craving for sweets, thirst, and sleepiness showed minimal change across this interval and no clear condition-dependent effects, suggesting that immediate post-ingestion shifts in these subjective states were limited.

### 3.3 Executive function

Executive function outcomes across post-ingestion time points (0–3 h) are shown in Figure 4. For flanker interference (reaction time difference between incongruent and congruent trials), a 2-way RM ANOVA revealed a significant main effect of condition, *F*(1, 14) = 5.400, *p* = 0.0452, η²p = 0.375, whereas the main effect of time, *F*(3, 42) = 1.471, *p* = 0.2445, η²p = 0.140, and the time × condition interaction, *F*(3, 42) = 1.161, *p* = 0.1666, η²p = 0.169, were not significant. Across the 0–3 h period, flanker interference was lower in the milk protein condition than in the control condition (predicted mean difference = 9.80 ms, 95% CI [0.26, 19.34]) (Fig. 4A).

**Figure 4.**
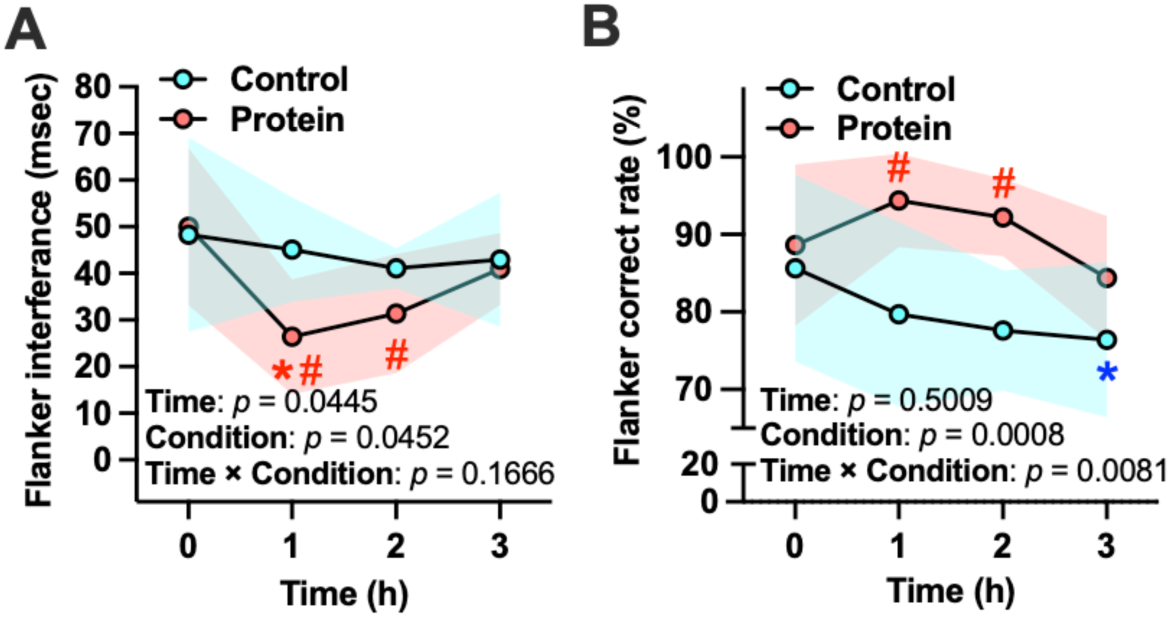
Executive function during prolonged esports play. Flanker task outcomes across post-ingestion time points (0–3 h) under the milk protein and control conditions. (A) Flanker interference (reaction time difference: incongruent − congruent). (B) Accuracy on incongruent trials (%). Data are shown as mean ± SD. The shaded bands represent the SD for the control and protein conditions. Results of the condition × time repeated-measures ANOVA are shown in the lower-left corner of each graph (Time, Condition, and Interaction). Where indicated, post-hoc comparisons were performed with Holm–Bonferroni correction. \**p* < 0.05 vs. 0 h within the same condition. #*p* < 0.05 vs. control condition at the indicated time point.

For incongruent-trial accuracy, a mixed-effects model revealed a significant main effect of condition, *F*(1, 14) = 20.70, *p* = 0.0008, η²p = 0.653, as well as a significant time × condition interaction, *F*(3, 42) = 5.474, *p* = 0.0081, η²p = 0.491. The main effect of time was not significant, *F*(3, 42) = 0.8035, *p* = 0.5009, η²p = 0.068. Overall, accuracy on incongruent trials was higher in the milk protein condition than in the control condition across the post-ingestion period (predicted mean difference = 9.34%, 95% CI [4.82, 13.85]) (Fig. 4B).

In supplementary analyses focusing on the baseline-to-immediate post-ingestion interval (Pre-intake to 0 h), flanker interference and incongruent-trial accuracy showed minimal change over time and no clear condition-dependent effects (5ementary Fig. 4). These results suggest that the executive-function differences observed during prolonged play primarily emerged during the gameplay period rather than immediately after ingestion.

To further link executive function outcomes to physiological indices, exploratory Pearson correlation analyses were conducted between pupil diameter and flanker performance using data from the 1–2 h gameplay interval (Fig. 5). Pupil diameter was negatively correlated with flanker interference time (r = −0.4505, 95% CI [−0.6732, −0.1529], p = 0.0045; Fig. 5A-C), whereas correlations with incongruent-trial accuracy were weaker and less consistent, reaching significance in only one analysis (r = 0.2887, 95% CI [−0.0090, 0.5393], p = 0.0474; Fig. 5D-F). As a robustness check, we repeated the correlation analyses using pupil and flanker values aggregated across 1–3 h (Supplementary Fig. 6).

**Figure 5.**
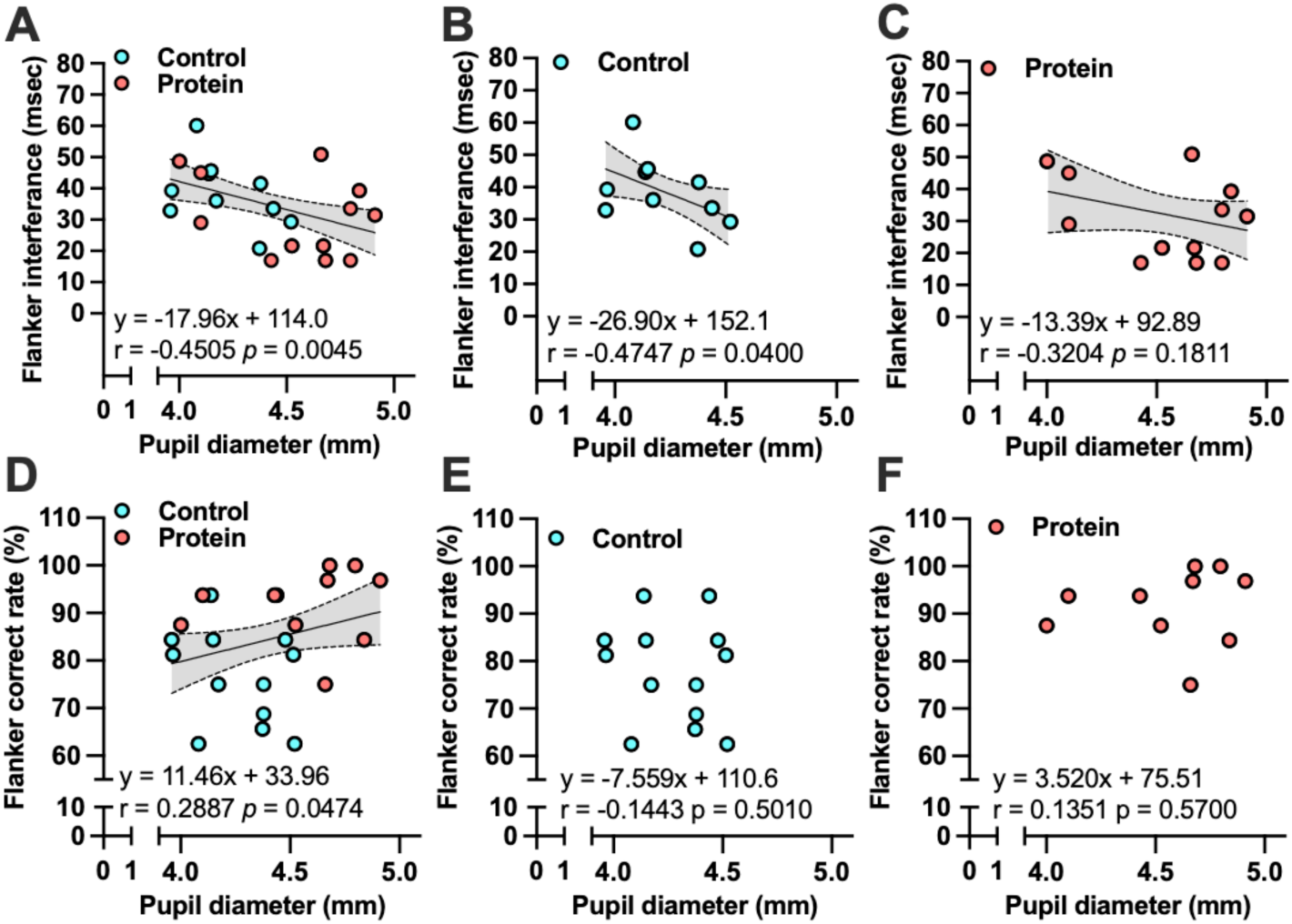
Associations between pupil diameter and flanker performance during early gameplay. Scatter plots show Pearson correlations between pupil diameter (mm) and flanker outcomes during the 1–2 h gameplay interval. (A) All participants (combined) for flanker interference time. (B) Control condition for flanker interference time. (C) Milk protein condition for flanker interference time. (D) All participants (combined) for incongruent-trial accuracy. (E) Control condition for incongruent-trial accuracy. (F) Milk protein condition for incongruent-trial accuracy. Correlation coefficients (r) and two-tailed p values are shown in each panel. The line and shaded band represent the fitted association and 95% confidence interval. Robustness-check correlations computed over the broader 1–3 h window is provided in Supplementary figure 5.

### 3.4 In-game performance metrics

In-game performance metrics during gameplay (1–3 h) are shown in Figure 6. Match-level statistics were averaged within each 1-h gameplay block to yield participant-level hourly performance measures.

**Figure 6.**
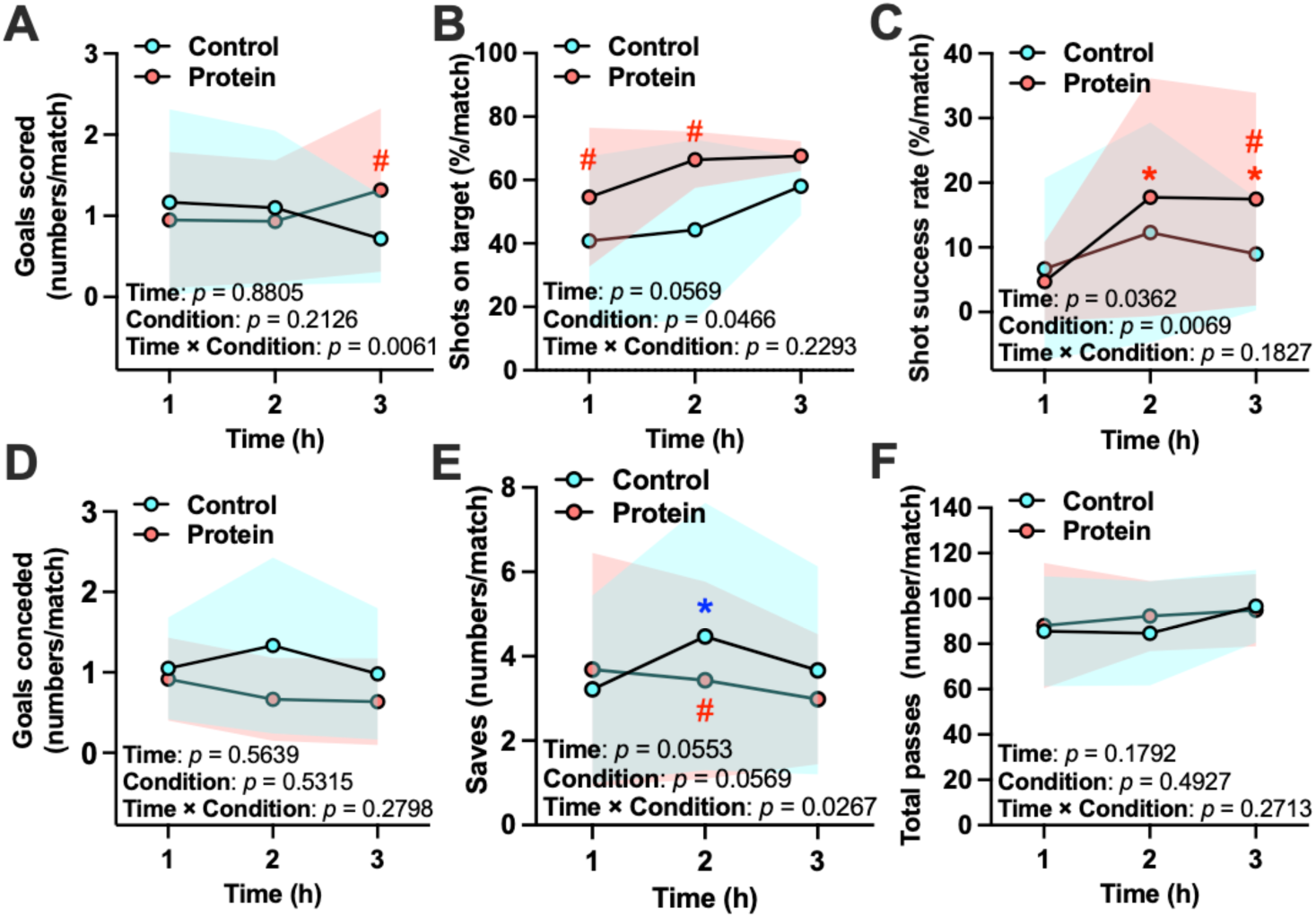
In-game performance metrics during gameplay. In-game performance metrics averaged within each 1-h gameplay block (1 h, 2 h, 3 h) under the milk protein and control conditions. Offensive metrics include (A) goals scored, (B) shots on target, and (C) shot success rate; defensive metrics include (D) goals conceded and (E) saves; (F) total passes are shown as an intermediate measure. Data are shown as mean ± SD. Results of the condition × time repeated-measures model are shown in the lower-left corner of each graph (Time, Condition, and Interaction). Where indicated, post-hoc comparisons between conditions at each time point were performed with Holm–Bonferroni correction. \**p* < 0.05 vs. 1st 1-h gameplay block (1 h) within the same condition. #*p* < 0.05 vs. control condition at the indicated time point.

#### Offensive performance

For goals scored, a significant time × condition interaction was observed, *F*(2, 28) = 6.868, *p* = 0.0061, η²p = 0.433, whereas the main effects of time, *F*(2, 28) = 0.128, *p* = 0.8805, η²p = 0.014, and condition, *F*(1, 14) = 1.800, *p* = 0.2126, η²p = 0.167, were not significant (Fig. 6A). For shots on target, no significant main effects or interaction were detected (time: *F*(2, 28) = 3.375, *p* = 0.0569, η²p = 0.273; condition: *F*(1, 14) = 3.441, *p* = 0.0966, η²p = 0.277; time × condition: *F*(2, 28) = 1.712, *p* = 0.2293, η²p = 0.255) (Fig. 6B). For shot success rate, significant main effects of time, *F*(2, 28) = 4.015, *p* = 0.0362, η²p = 0.309, and condition, *F*(1, 14) = 12.11, *p* = 0.0069, η²p = 0.574, were observed, whereas the time × condition interaction was not significant, *F*(2, 28) = 1.871, *p* = 0.1827, η²p = 0.172 (Fig. 6C).

#### Defensive performance

For goals conceded, no significant main effects or interaction were observed (time: *F*(2, 28) = 0.592, *p* = 0.5639, η²p = 0.062; condition: *F*(1, 14) = 0.423, *p* = 0.5315, η²p = 0.045; time × condition: *F*(2, 28) = 1.587, *p* = 0.2798, η²p = 0.346) (Fig. 6D). For saves, a significant time × condition interaction was detected, *F*(2, 28) = 4.463, *p* = 0.0267, η²p = 0.331, whereas the main effects of time, *F*(2, 28) = 3.416, *p* = 0.0553, η²p = 0.275, and condition, *F*(1, 14) = 4.764, *p* = 0.0569, η²p = 0.346, did not reach statistical significance (Fig. 6E).

#### Intermediateh performance measure

For total passes, no significant main effects or interaction were observed (time: *F*(2, 28) = 1.895, *p* = 0.1792, η²p = 0.174; condition: *F*(1, 14) = 0.511, *p* = 0.4927, η²p = 0.054; time × condition: *F*(2, 28) = 1.404, *p* = 0.2713, η²p = 0.135) (Fig. 6F).

## 4. Discussion

In this randomized, single-blind, energy-matched controlled crossover study, we examined whether acute milk protein intake attenuates fatigue-related physiological responses during prolonged esports play and supports subjective state, executive control, and in-game performance (Fig. 1). Overall, milk protein intake was associated with convergent physiological changes, including larger pupil diameter during gameplay, smoother interstitial glucose dynamics, and lower salivary cortisol (Fig. 2). These physiological changes were accompanied by higher enjoyment and lower hunger (Fig. 3) and improved executive function on the flanker task (Fig. 4). Consistent with a physiological–cognitive coupling, larger pupil diameter during gameplay was associated with better inhibitory control on the flanker task (Fig. 5). Finally, milk protein intake was associated with condition-dependent changes in in-game offensive performance metrics, most notably a higher shot success rate (Fig. 6). Together, these findings support the proposed physiological-to-behavioral framework in which acute milk protein intake helps sustain esports performance during prolonged play, consistent with milk protein–based smart nutrition as a practical, non-stimulant approach designed to sustain cognitive–behavioral performance while minimizing health trade-offs.

### Protein-supported arousal and inhibitory control during prolonged esports play

In the present study, milk protein intake was accompanied by larger pupil diameter during gameplay, consistent with attenuation of fatigue-associated pupil constriction (Fig. 2A). Mechanistically, pupillary dynamics provide a useful physiological window into the state that supports inhibitory control during prolonged esports play. Pupil diameter is commonly used as a noninvasive marker of central arousal and cognitive resource allocation and has been linked, albeit imperfectly, to neuromodulatory mechanisms relevant to executive control, including locus coeruleus–noradrenergic function (Allen, 2020; Aston-Jones & Cohen, 2005).

A plausible nutritional pathway is that dietary protein intake alters circulating amino acid profiles, thereby modulating the availability of precursor amino acids for neurotransmitters and neuromodulators relevant to cognitive control, including catecholamines (Fernstrom & Fernstrom, 2007; Lieberman, 2003). In cognitively demanding contexts, such precursor availability, particularly for aromatic amino acids such as tyrosine, has been proposed to become functionally relevant for performance (Jongkees et al., 2015a).

Linking physiology to cognition, pupil diameter during early gameplay was negatively associated with flanker interference time, such that smaller pupil diameters co-occurred with larger interference costs, whereas associations with incongruent-trial accuracy were weaker and less consistent (Fig. 5). This pattern is compatible with evidence that pupillary measures are often more sensitive to changes in cognitive effort and response efficiency during control-demanding tasks than to accuracy per se, which can be subject to ceiling effects (van der Wel & van Steenbergen, 2018). Although correlational, these findings provide convergent support for a physiological–cognitive coupling in which maintaining central arousal during prolonged play is related to better inhibitory control.

### Protein-stabilized glycemic dynamics and appetite during prolonged esports play

A second mechanistic pathway supported by our data involves metabolic dynamics and appetite-related subjective state. Although time-averaged interstitial glucose did not differ between conditions, milk protein intake produced a clearly different temporal glucose profile during the post-ingestion period, indicating smoother glycemic dynamics across prolonged play, consistent with reduced glycaemic variability over time (Fig. 2C).

Mechanistically, milk-derived proteins (including whey-containing products) can blunt postprandial glycaemic excursions by slowing gastric emptying and enhancing insulin and incretin responses (e.g., GLP-1), thereby moderating the rate of glucose appearance in the circulation (Jakubowicz et al., 2014; Nesti et al., 2019). Such temporal glucose features are behaviorally relevant because postprandial glucose “dips” and glycemic instability have been linked to subsequent hunger and energy intake in healthy individuals (Wyatt et al., 2021). This point may be particularly relevant in esports and other prolonged digital tasks, where players often seek alertness from sugar-containing beverages that can provide transient benefit yet may also promote glycaemic variability that is less compatible with sustained cognitive–behavioral stability over time.

Consistent with this framework, participants reported lower hunger ratings and craving for sweets in the milk protein condition across 0–3 h, with a time-dependent condition difference (Fig. 3 & Supplementary Fig. 3). Together, these findings suggest that acute milk protein intake may help maintain a more stable metabolic milieu during prolonged esports play, which in turn could reduce appetite-related distraction and support sustained engagement. These acute post-ingestion effects are also evident in the baseline-to-0 h interval (Supplementary Fig. 2).

### Protein-related enhancement of soccer-like performance

A third implication of our framework concerns behavioral translation in soccer-like performance metrics. Milk protein intake was associated with condition-dependent improvements in in-game performance during prolonged virtual soccer play (Fig. 6), most notably a higher shot success rate across gameplay blocks and time-dependent effects on goals scored (Fig. 6C). Soccer performance depends not only on physical execution but also on sustained cognitive control and efficient decision-making under time pressure; accordingly, experimental and review evidence indicates that cognitive/mental fatigue can impair soccer-relevant technical skills and decision-making, which can reduce efficiency in key actions such as shooting (Sun et al., 2022; Trecroci et al., 2020).

In the present esports context, the behavioral benefits observed in virtual soccer co-occurred with improved executive control (Fig. 4) and a more favorable physiological profile during gameplay (Figs. 2 and 3), consistent with the interpretation that acute milk protein intake helped preserve cognitive resources that support performance efficiency under fatigue (Matsui et al., 2024). This interpretation is also consistent with prior clinical evidence that whey peptide supplementation can improve cognitive performance, particularly in individuals reporting higher subjective fatigue (Kita et al., 2018).

Mechanistically, such an effect is plausible given evidence that catecholamine precursor availability (e.g., tyrosine-related pathways) can become performance-relevant under cognitively demanding or stressful conditions, supporting control processes that contribute to efficient action selection (Jongkees et al., 2015b). Notably, total passes did not differ between conditions (Fig. 6F), suggesting that the intervention did not simply alter overall play involvement or strategy, but rather the effectiveness of goal-directed actions.

On the defensive side, the time-dependent effect on saves (Fig. 6E) further suggests that fatigue-sensitive control processes may extend to defensive performance, although goals conceded did not show a clear condition effect. Because match events can still be influenced by situational dynamics even under standardized offline settings, these in-game outcomes should be interpreted as behavioral readouts that align with the physiological and cognitive findings rather than as deterministic indicators of overall skill.

### Broader implications for prolonged digital cognitive work

Esports represents an increasingly prevalent form of sustained, high-load digital cognition that combines prolonged screen exposure, continuous decision-making, and time-pressured executive control. Similar cognitive fatigue processes and performance costs are also reported in other prolonged digital or knowledge-work contexts, where sustained mental effort can impair decision-making and self-regulation (Brady et al., 2024). In parallel, pupillometry and related ocular metrics are increasingly used in human–computer interaction and applied settings to index cognitive workload and engagement over time, supporting the relevance of the present physiological markers beyond esports (Gorin et al., 2024). From this perspective, the coherent pattern observed here across physiology, subjective state, and executive control suggests that milk protein–based nutrition could be explored as a practical, non-stimulant strategy that supports sustained cognitive–behavioral performance while minimizing health trade-offs, in other prolonged digital activities (e.g., extended computer-based work or training) that share key fatigue demands with esports.

### Limitations and future directions

Several limitations should be considered when interpreting these findings. First, the sample size was modest and comprised healthy young adult men with esports experience, which may limit generalizability to broader player populations, including elite esports athletes, women, older adults, and individuals with metabolic or sleep-related conditions. Second, although the crossover design and standardized game settings strengthen internal validity, in-game performance metrics can still be influenced by situational dynamics, and the present results were obtained in offline matches against a computer-controlled opponent; replication across game genres, competitive modes, and ecologically richer contexts is warranted. T Third, although beverages were served in identical opaque containers, sensory differences between the milk protein drink and apple juice may have partially unblinded participants and influenced subjective outcomes and performance; accordingly, the study is more appropriately characterized as single-blind (assessor-blind) rather than double-blind. Fourth, mechanistic conclusions remain tentative because neurotransmitter activity and related metabolic mediators (e.g., insulin/incretin responses) were not directly measured; future studies incorporating targeted biomarkers and causal designs will be needed to test the proposed pathways. Fifth, all sessions were conducted in the morning (08:30–09:00 start), and potential time-of-day effects on physiological and cognitive responses mean that the generalizability of these findings to afternoon or evening intake remains unknown. Despite these limitations, the present findings provide evidence that an acute milk protein drink may help sustain physiological state and cognitive–behavioral performance during prolonged virtual soccer play, supporting the potential of smart nutrition as a practical, non-stimulant strategy for esports.

## 5. Conclusion

This randomized, single-blind, energy-matched controlled crossover study indicates that acute milk protein intake can help sustain performance during prolonged esports (virtual soccer) play. Milk protein intake was associated with larger pupil diameter during gameplay, smoother interstitial glucose dynamics, and lower salivary cortisol, together with higher enjoyment and lower hunger, improved flanker performance, and better in-game efficiency, most notably higher shot success rate. These findings support milk protein–based smart nutrition as a practical, non-stimulant, short-term nutritional approach to sustaining cognitive–behavioral performance during prolonged digital cognitive activities such as esports.

## Data availability

The data supporting the findings of this study are available from the corresponding author upon request.

## Author contribution

T.M., C.O. and K.N. conceived and designed the study. T.M., S.T., and D.F. recruited participants, collected the data, and conducted data analysis. T.M., S.T., D.F., C.O., and K.N. interpreted the data. T. M. drafted the manuscript. T.M., S.T., C.O. and K.N. edited and revised the manuscript. All authors approved the final version.

---

**Supplementary figure 1.**
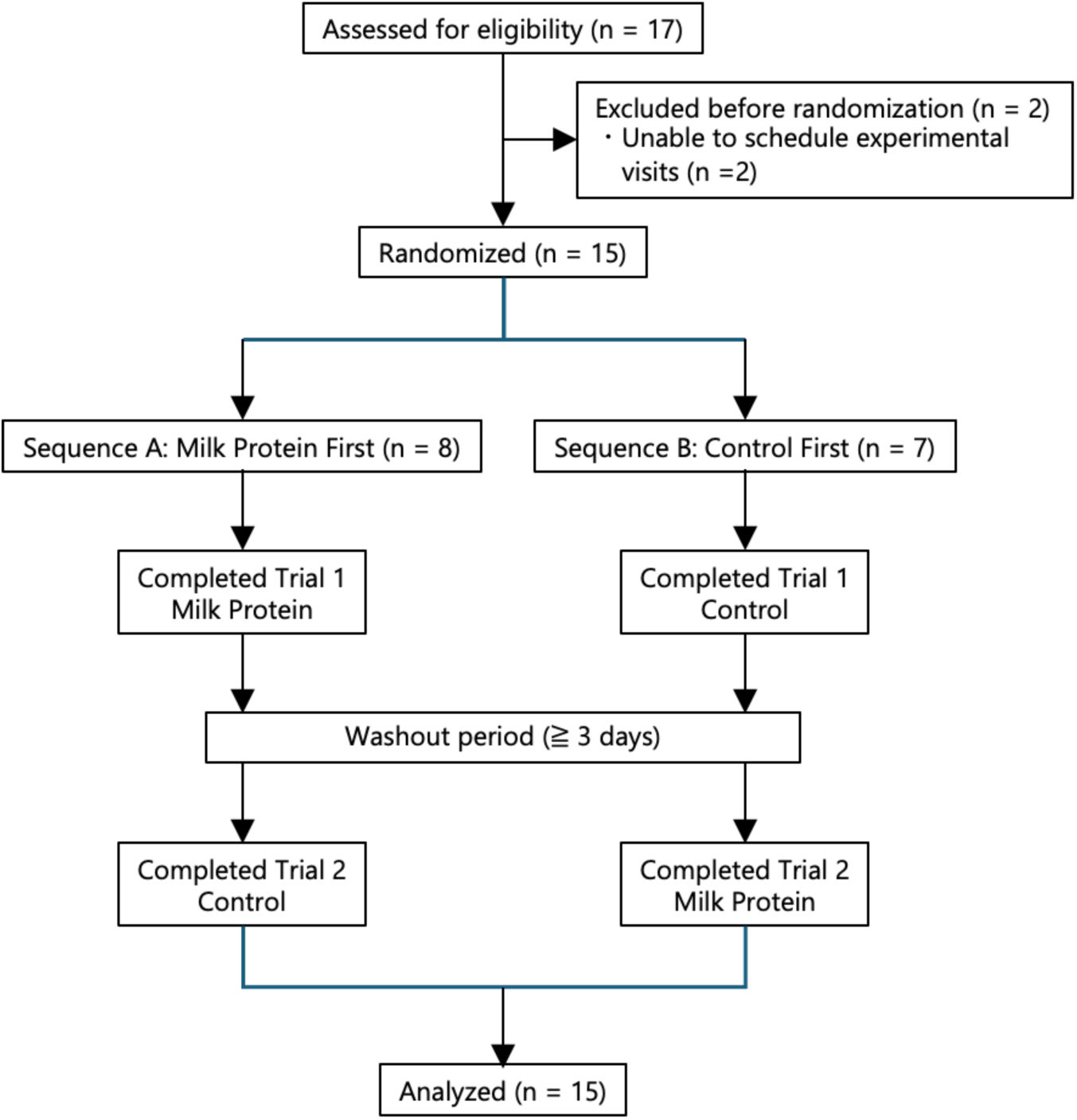
Participant flow diagram. Flow diagram of participant enrollment, randomization, crossover intervention, and analysis. A total of 17 individuals were assessed for eligibility, of whom 2 withdrew before randomization because the experimental visits could not be scheduled. The remaining 15 participants were randomized to one of two sequences: milk protein followed by control (Sequence A, n = 8) or control followed by milk protein (Sequence B, n = 7). All randomized participants completed both experimental sessions, which were separated by a washout period of at least 3 days, and were included in the final analysis.

**Supplementary figure 2.**
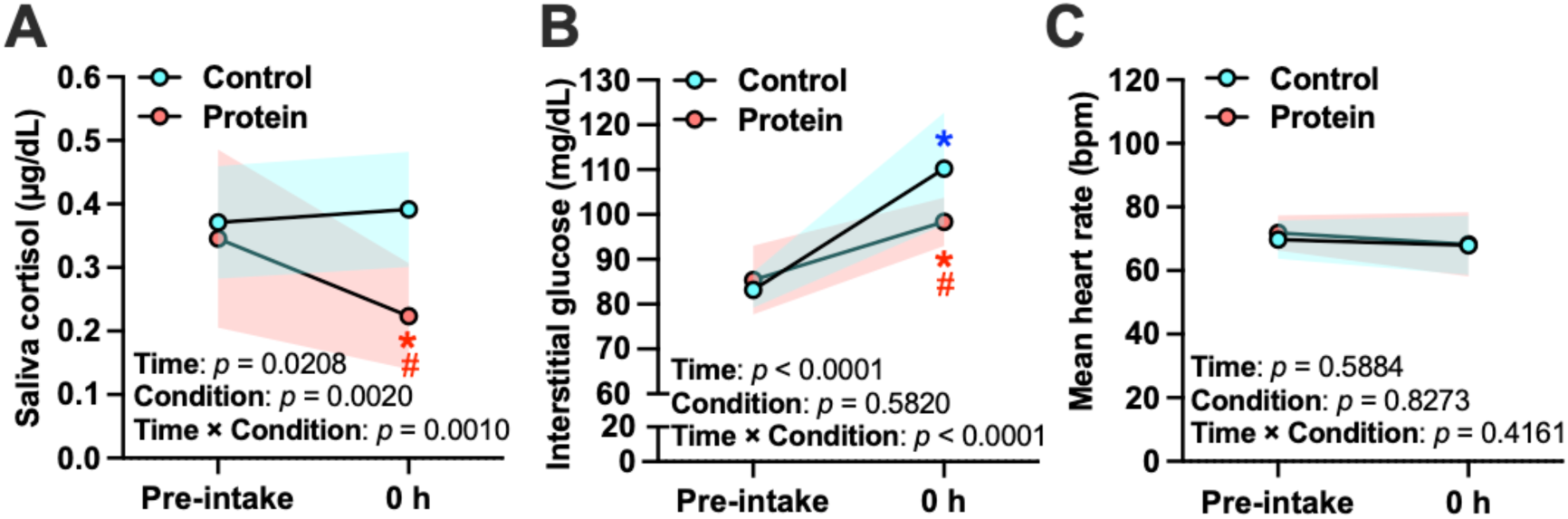
Baseline and immediate post-ingestion physiological responses. Baseline (Pre-intake) and immediate post-ingestion (0 h) values for (A) salivary cortisol, (B) interstitial glucose measured by continuous glucose monitoring (CGM), and (C) mean heart rate under the milk protein and control conditions. Data are shown as mean ± SD. Shaded bands indicate the SD for the control and protein conditions. Condition × time repeated-measures ANOVA results are shown within each panel (Time, Condition, and Interaction). Where indicated, Holm–Bonferroni–corrected post-hoc tests were used for within-condition comparisons between time points. *p < 0.05 vs. Pre-intake within the same condition; #p < 0.05 vs. the control condition at the indicated time point (Holm–Bonferroni corrected).

**Supplementary figure 3.**
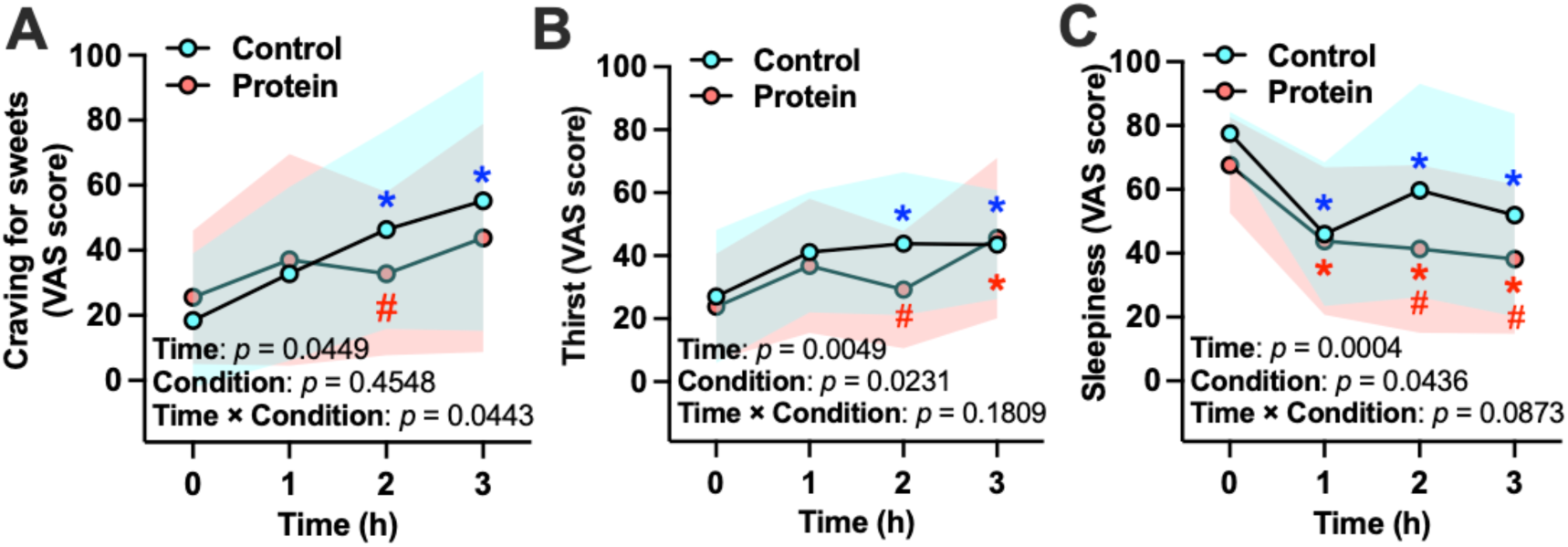
Secondary subjective ratings during prolonged esports play. Secondary visual analogue scale (VAS) ratings for (A) craving for sweets, (B) thirst, and (C) sleepiness measured during prolonged esports play (0–3 h) under the milk protein and control conditions. Data are shown as mean ± SD. Shaded bands indicate the SD for the control and protein conditions. Condition × time repeated-measures ANOVA results are shown within each panel (Time, Condition, and Interaction). Where indicated, Holm–Bonferroni–corrected post-hoc tests were used for within-condition comparisons between time points. *p < 0.05 vs. 0 h within the same condition; #p < 0.05 vs. the control condition at the indicated time point (Holm–Bonferroni corrected).

**Supplementary figure 4.**
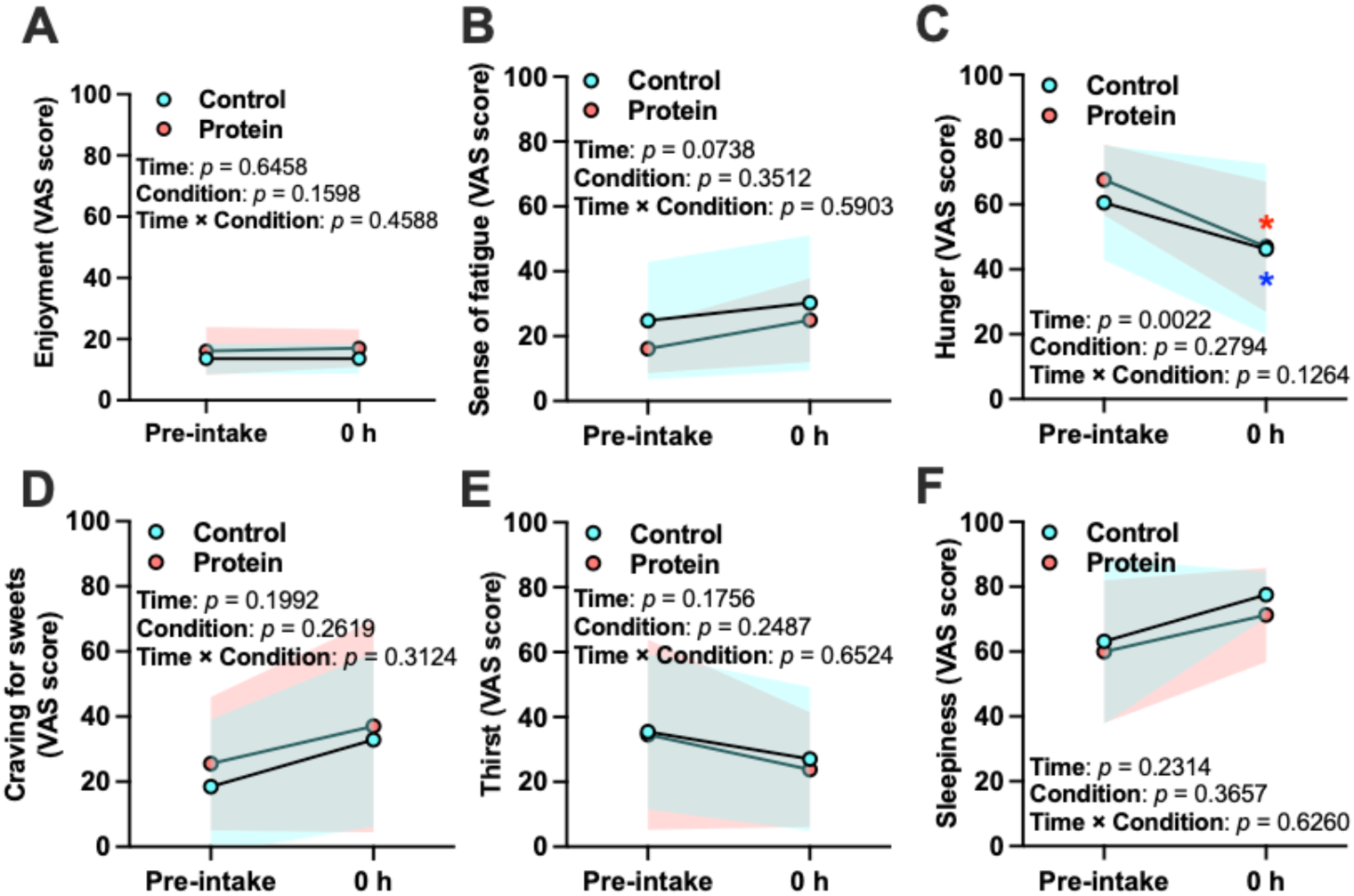
Baseline and immediate post-ingestion subjective ratings. Baseline (Pre-intake) and immediate post-ingestion (0 h) visual analogue scale (VAS) ratings for (A) enjoyment, (B) sense of fatigue, (C) hunger, (D) craving for sweets, (E) thirst, and (F) sleepiness under the milk protein and control conditions. Data are shown as mean ± SD. Shaded bands indicate the SD for the control and protein conditions. Condition × time repeated-measures ANOVA results are shown within each panel (Time, Condition, and Interaction). Where indicated, Holm–Bonferroni–corrected post-hoc tests were used for within-condition comparisons between time points. *p < 0.05 vs. Pre-intake within the same condition (Holm–Bonferroni corrected).

**Supplementary figure 5.**
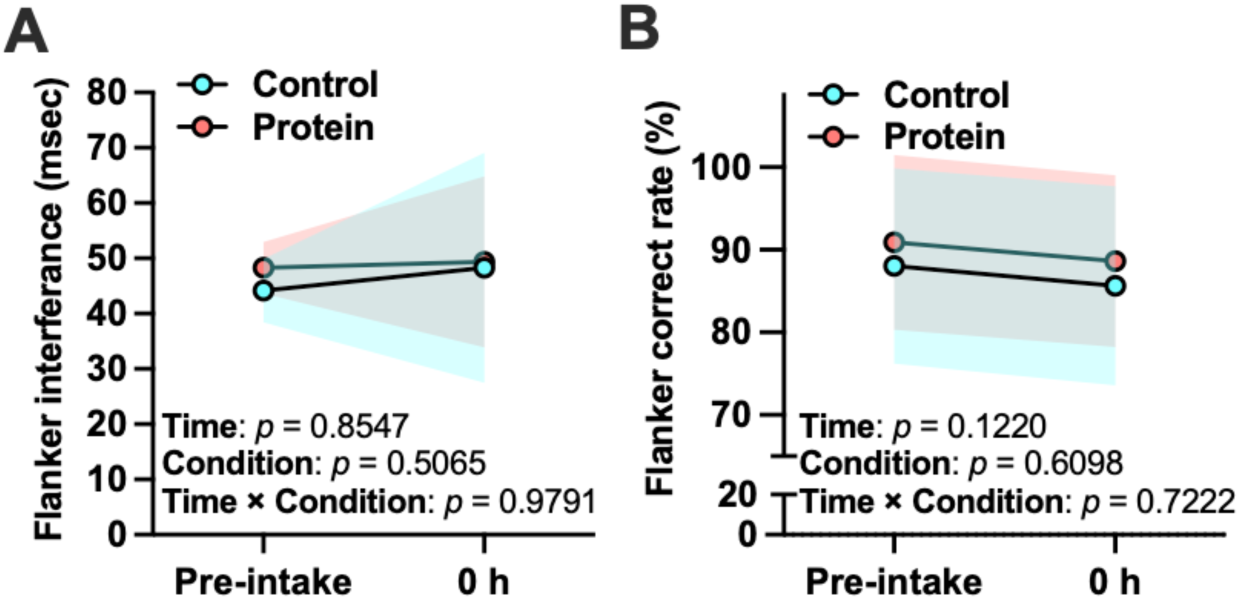
Baseline and immediate post-ingestion executive function. Baseline (Pre-intake) and immediate post-ingestion (0 h) flanker task outcomes under the milk protein and control conditions. (A) Flanker interference (reaction time difference: incongruent − congruent). (B) Accuracy on incongruent trials (%). Data are shown as mean ± SD. Shaded bands indicate the SD for the control and protein conditions. Condition × time repeated-measures ANOVA results are shown within each panel (Time, Condition, and Interaction).

**Supplementary figure 6.**
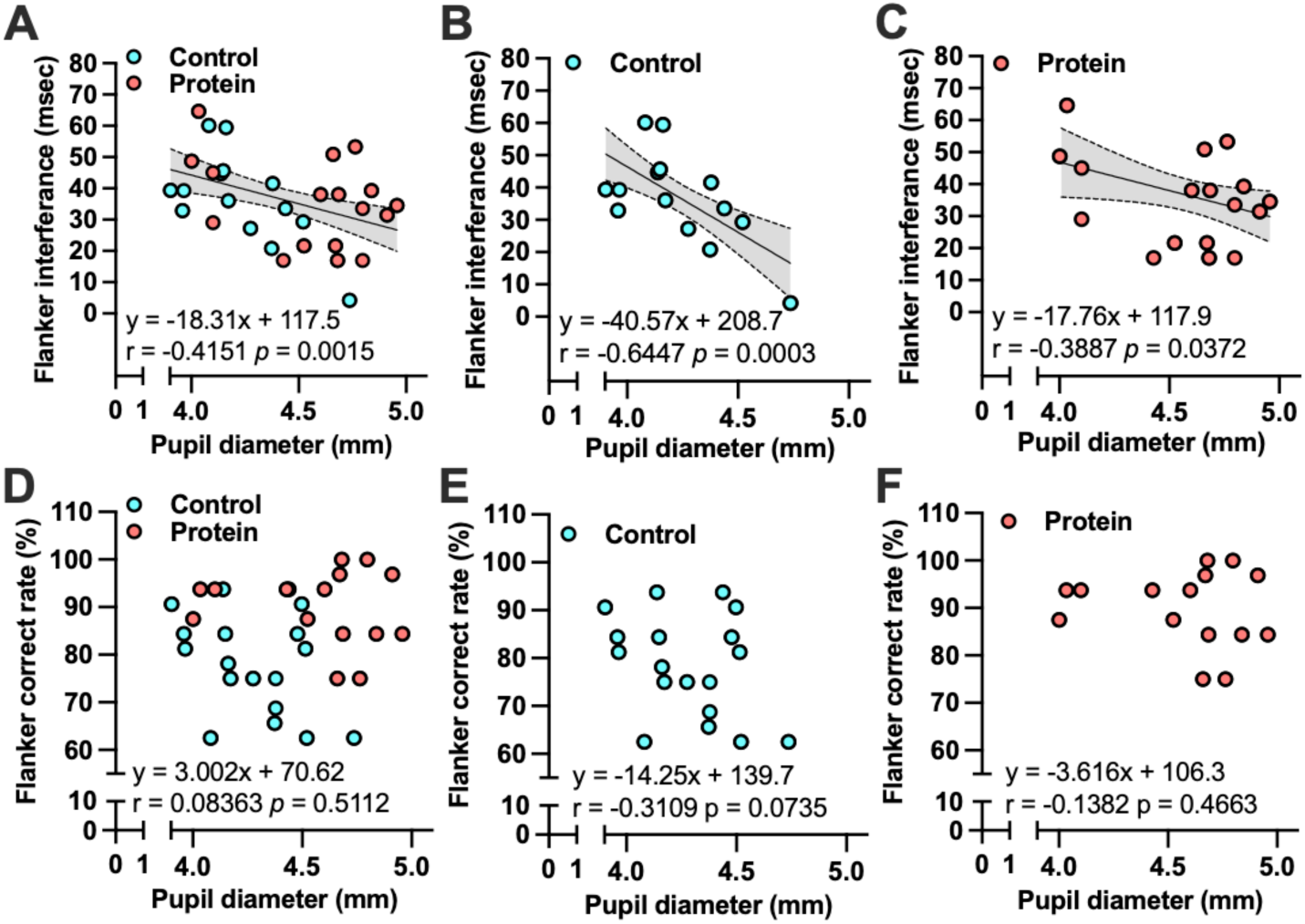
Robustness-check correlations between pupil diameter and flanker performance across gameplay. Scatter plots show Pearson correlations between pupil diameter (mm) and flanker outcomes computed over the broader gameplay window (1–3 h) as a robustness check. (A) All participants (combined) for flanker interference time. (B) Control condition for flanker interference time. (C) Milk protein condition for flanker interference time. (D) All participants (combined) for incongruent-trial accuracy. (E) Control condition for incongruent-trial accuracy. (F) Milk protein condition for incongruent-trial accuracy. Solid lines indicate linear regression fits with 95% confidence bands. Correlation coefficients (r) and two-tailed p values are shown in each panel. The line and shaded band represent the fitted association and 95% confidence interval.

